# Embryonic life histories in annual killifish: adapted to what?

**DOI:** 10.1101/2023.08.03.551794

**Authors:** Tom JM Van Dooren

**Affiliations:** team VPA Phenotypic Variability and Adaptation, CNRS, Institute for Ecology and Environmental Sciences iEES Paris, Sorbonne University, 4, Place Jussieu, 75005 Paris France

**Keywords:** Adaptation, environmental change, annual killifish, diapause, pond life cycle, eco-evo-devo, cumulative hazards, longitudinal studies, Adaptive Dynamics

## Abstract

Adaptation requires an evolving strategy and an environment. Given an environment, we predict or estimate which strategies are adapted. Given a strategy, we want to know in which environments it might be adapted. Example calculations aiming to determine such environments, named evolutionarily singular environments ESE, are presented using lab data on embryonic life histories of *Austrolebias* annual killifish. Annual killifish embryos can arrest development and survive desiccation of temporary ponds in the soil. They might implement diversified bet-hedging, generally seen as an adaptation to uncertain environments. Using parameter estimates and parsimonious assumptions, a population dynamical model is constructed with explicit developmental stages. Using invasion fitness gradients of rates of development and hatching probabilities, it is investigated whether these could be adapted to pond filling regimes with gradual filling and drying and deterministic within-year variation only. The life history as a whole is not adapted to the regular within-year annual cycles investigated, with one or two periods where reproduction can occur. Faster development rates were always favoured, just as in constant environments. Only for hatching probabilities, pond filling regimes were found which made their invasion fitness sensitivities zero. However, the observed trait values did not have long-term evolutionary stability in these ESE. Therefore, neither the developmental rates nor the hatching strategy seem adapted to within-year patterns of environmental change.

## Introduction

Evolutionary adaptation requires traits which can evolve and an environment. Part of the environment, the “external driver” (e.g., Metz et al. 2008), is usually seen as given and not affected by changes in an evolving population. Effects of environmental variation on adaptation are assessed by the population structuring they generate (Southwood 1977), and through the means and variances of demographic parameters (Cohen 1966, Simons 2011, Starrfelt and Kokko 2012). In modelling, the actual environment often figures by means of examples congruent with assumptions on demographic parameters (e.g., ten Brink et al. 2020). Similarly, calls for better predictions of adaptation to climate change urge more elaborate demographic and trait models (Urban et al. 2023) but it is unclear whether we are grasping the environments themselves and their effects on demography well. In evolutionary conservation biology, adaptive population management (e.g., Zhan et al. 2015) includes controlling the environment, which then isn’t external nor a given.

Evolutionary anthropologists have discussed the “environment of evolutionary adaptedness” (Bowlby 1973), the environment in which a population evolved and presumably became adapted. Historical aspects of this definition were problematic (Irons 1998). We currently favour a rather a-historical view of adaptation (Reeve & Sherman 1993), where an adapted suite of traits confers highest fitness among a set of alternatives in a given environment. For a given organismal strategy, observed in a single or across different environments, we often want to know in which environments it would be adapted. It would be the most common strategy in them (Reeve & Sherman 1993) or one of the strategies maintained in a protected polymorphism (Geritz et al. 1998). In these environments, there is no long-term directional selection giving rare alternative strategies a systematic advantage (e.g., Ellner 1997, Coulson 2021). For adaptive management, conservation, and to investigate whether we have understood the environment sufficiently, knowing such environments seems useful.

Here, an example is presented of a search for environments to which an observed strategy could be adapted. This is done for embryonic life history traits in an annual killifish (Van Dooren & Varela-Lasheras 2018). Annual killifish are oviparous cyprinodontiform fish species from the suborder Aplocheiloidei which evolved “annualism” repeatedly (Furness et al. 2015a, Helmstetter et al. 2016, Thompson et al. 2021). They use embryonic diapauses to persist in temporary ponds in Africa and South America which dry out seasonally (Berois et al. 2015) and where the start and end of a wet period generally vary between years. Many annual fish species are threatened (Volcan & Lanés 2018). Their habitats are restricted in space and managing them seems feasible. We could aim to keep annual fish populations adapted by preserving or changing their environments, not by seeing these as given.

Data patterns in annual killifish embryonic life histories have been interpreted as in agreement with bet-hedging adaptations to environmental uncertainty (Furness et al. 2015b, Pinceel et al. 2015, Polačik et al. 2017, Van Dooren & Varela-Lasheras 2018), but also to predictable properties of the environments. Furness et al. (2015b) proposed that a diapause occurs less or not at high temperatures because these predict the probability that another generation can be completed before the cool dry season starts. Similarly, Polačik et al. (2017) proposed that eggs from younger mothers develop faster to be ready when an extra opportunity to hatch and reproduce within a season presents itself. However, in *Austrolebias* annual fish (Alonso et al. 2023) for example, embryos will not hatch and grow when they have not experienced a dry period yet. The environment proposed where this would be adaptive therefore needs to be an environmental regime with two periods within a year where reproduction can occur, separated by at least partial drying of a pond.

Complete drying would avoid competition between cohorts hatched in the two periods. A demographic model and fitness calculations across the entire life cycle (Childs et al. 2010, de Jong et al. 2011, Van Dooren & Varela-Lasheras 2018) in different seasonal patterns of wet and dry periods are required to investigate whether the observed patterns would be adapted to them.

Such a model is presented here and it is used to investigate whether developmental rates and hatching probabilities might be adaptations to deterministic within-year environmental patterns. The model makes two important assumptions: it is assumed that a developmental strategy can only be adapted when there is no persistent directional selection on it. Second, data observed on individual development in lab conditions are assumed to be representative of what individuals do in the field. The model structure is tailored to specifics of annual killifish life histories and is integrating estimated parameters mostly from *A. bellottii*, the Argentinian pearl killifish (Steindachner 1881, Van Dooren & Varela-Lasheras 2018). The results demonstrate that the traits of observed embryonic life histories are not adapted to environments changing deterministically within years. Faster developmental rates are always favoured. For hatching probabilities, however, directional selection vanishes in some environments investigated. Based on the results, expectations are formulated for the adaptiveness of patterns when between-year variation occurs, deterministic or not.

## Methods

In the model detailed below, annual fish individuals can find themselves in different pond compartments and environments (Figure one). Environmental and population states are projected forward in time on a day-to-day basis.

**Figure one.**
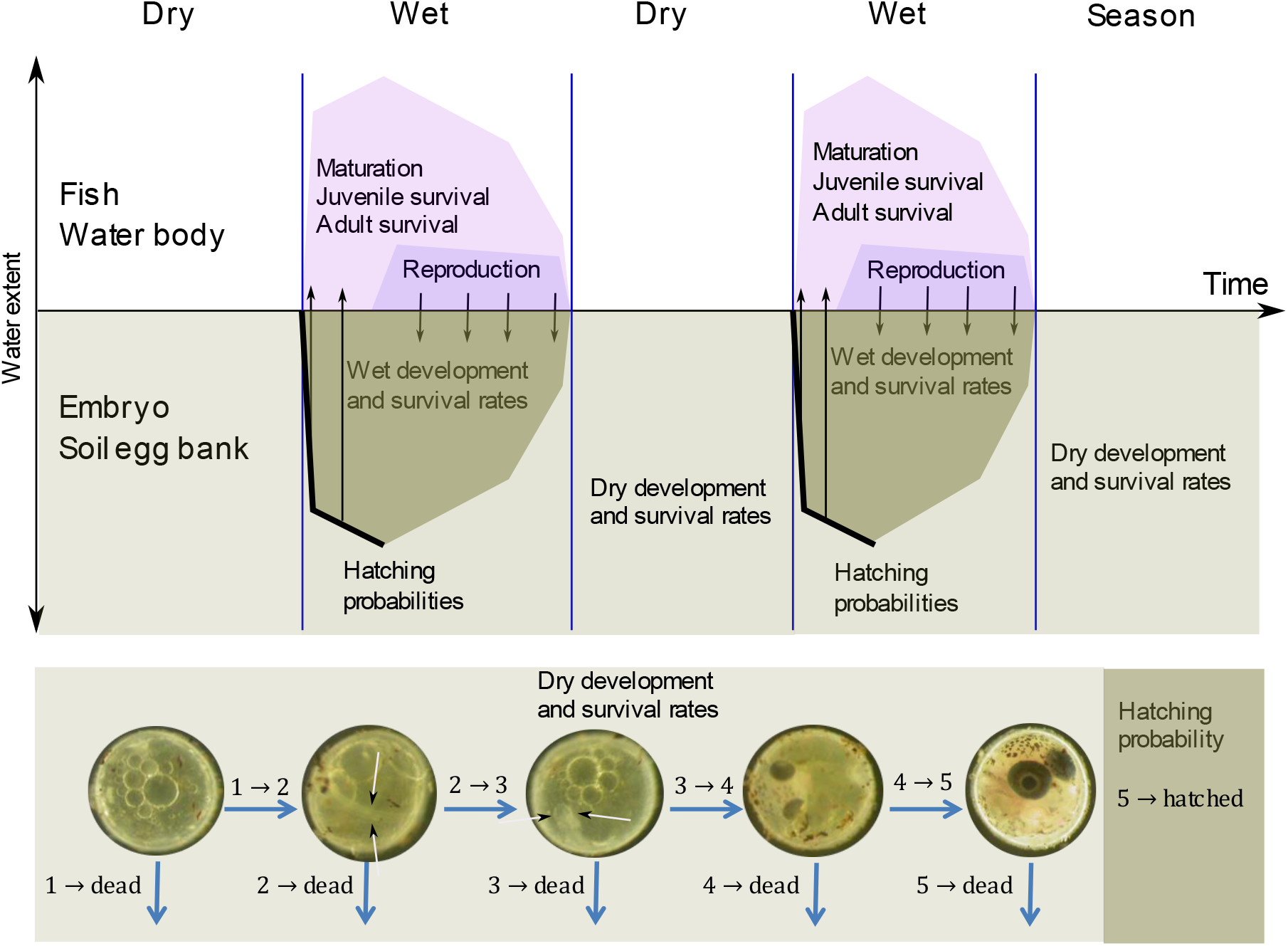
Schematic depiction of the annual killifish life history interacting with the pond environment. Above. Main processes over time in water and soil compartments and exchanges between them are listed. The egg bank is located in the soil compartment and fish in the water compartment. Dry and wet periods alternate: during a wet period at least part of the pond is filled (blue). Embryos in the soil egg bank cay be in dry (pale drab) or wet conditions (drab), depending on the extent of the pond covered by water. Individuals can hatch from stage five embryos when water reaches the location where they are buried (thick black line), survive and mature into adults. Reproducing adults contribute eggs to the soil egg bank at locations where water is still covering the soil. Below. A depiction of the developmental strategy modelled. Five developmental stages are considered, each with stage-specific mortality and development rates in dry conditions. Stage five embryos can hatch according a sub-model for hatching probabilities. Stages four and five are easy to discern due to their pigmented eyes. In stage two and three the body axis can be seen already, in stage three with a clear head (white arrows). Photographs of embryos reproduced from Van Dooren & Varela-Lasheras (2018) with permission.

### Embryonic life histories

The strategy for which we want to find environments where it would be adapted, consists of rates of development and hatching probabilities estimated for a group of embryos of the Argentinian pearl killifish *A. bellottii* (Van Dooren & Varela-Lasheras 2018). The data used to parameterize the embryonic life history were collected at 19.5C and 20.5C, while the average annual temperature for their site of origin was 17.9C (Ingeniero Maschwitz Argentina, Worldclim value used in Helmstetter et al. 2020). In this section, next to detailing the given strategy, we also include the description of mortality rates in different conditions, so that this section can be seen as the description of life history events in the egg bank compartment of the model. Events occur with different rates when embryos are in (1) wet conditions (e.g., their location covered by water and conditions hypoxic), when they are in (2) dry conditions (oxygenated with limited desiccation) and haven’t been rewetted yet (first dry period) and (3) in subsequent dry periods. In dry conditions, mortality rates differ between developmental stages.

The cohort from which development rates were estimated for “dry” conditions started incubation in wet oxygenated conditions. After some time, it was further incubated in air with 100% relative humidity (Supplementary Material). Five distinct developmental stages were considered (Figure one, Van Dooren & Varela-Lasheras 2018, simplified from Wourms 1972a), each with an age-dependent pattern of the risk to enter the next developmental stage in the first four developmental stages.

Embryos were assigned to stage one at age zero. In stage two, a neural keel and somites have appeared. In stage three, optic cups were visibly present. In stage four, the eyes had become pigmented. In the fifth stage the embryo completely surrounded the yolk sac. In the conditions where the strategy has been determined, diapause has been observed in *Austrolebias bellottii* in stages one (DI) and five (DIII, Van Dooren & Varela-Lasheras 2018) as age intervals with negligible risk to enter the next stage. From stage five, embryos can hatch when they are rewetted. Hatching probabilities were estimated using logistic regressions (Van Dooren & Varela-Lasheras 2018). It was noted that after some time, the rate of events (deaths or developmental transitions) decreased to almost zero. A similar observation was made by Polačik & Vrtilek (2023). This slowdown is more prominent even after the first rewetting of eggs and we incorporated that into the life history model in an ad-hoc manner as explained below.

For the first dry period embryos experience, estimated cumulative hazards were used to obtain transition probabilities between developmental stages or from any developmental stage to the dead state (Supplementary Material). Fifteen different transitions (from each developmental stage to all more advanced ones and to the dead state) can be observed across a time interval, some more often than others. It is only age given in the number of days in dry conditions which is assumed to affect developmental progress, therefore we use this integer as the individual age variable. Matrices were derived with transition probabilities per day using age-dependent multi-state modelling with competing risks (de Wreede et al. 2011). Van Dooren & Varela-Lasheras (2018) found that many embryos don’t resume development after a first rewetting. To integrate this in the model in a straightforward manner, a 1/0 state variable was added to the individual state denoting whether an embryo would be in a state where development proceeds (1), or remains in a generally arrested state (0). Each time an embryo experiences wet conditions after the first dry period, it enters this non-advancing state with probability *p*_d_ which is maintained during the next dry period. The probability of survival per day 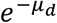 in the non-advancing state, which was equal in all stages, and the proportion entering the advancing state *p*_d_ were estimated based on the data in Van Dooren & Varela-Lasheras (2018) and are given in Table one.

When stage five embryos are rewetted (i.e., covered by standing water again), they can hatch on that day only in the model. This occurs with probabilities which differ between the first and later rewettings and with an age effect at first rewetting (Van Dooren & Varela-Lasheras 2018). Three hatching parameters *β*_1_, *β* _2_, *β* _3_ required to model this were estimated and are given in Table one. Upon first rewetting, embryos in stage five hatch with probability *g*(*β*_1_ + *β*_2_*a*), with *g* the inverse logit function and *a* the number of days an embryo has been dry. First rewetting is therefore an 1/0 state variable. At later rewettings, embryos hatch with probability *g*(*β* _3_). Eggs which don’t hatch remain in the soil at the same location. In wet soil conditions, embryonic development does not proceed (Van Goor 2009, Polačik et al. 2021, 2023), presumably due to lack of oxygen. In a pilot study on the related species *Acantholebias luteoflammulatus* (van Goor 2009), individual survival was followed in the field. The probability per day that an egg survives in wet conditions 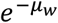 was computed from the results and used across all stages (Table one). The value used is lower than an estimate (0.997) based on the data in Polačik et al. (2023). There are no stage-specific survival probabilities in wet soil estimated yet. Fitness effects of developmental rates per stage in dry conditions and in the advancing state were predicted as detailed below, and fitness effects of hatching parameters *β*_1_ and *β* _3_. The probabilities of individual switching between 1/0 states and of the age plasticity of hatching were not treated as traits which can evolve. The parameter values for these transitions were based on observations at few censuses and considered too much ad-hoc in comparison to other embryonic life history parameters.

### Evolutionarily Singular Environments

Given a strategy, we want to know in which environments it might be adapted. When we start from a vector of scalar trait values ***x*** representing a strategy (for function-valued ***x***, see Dieckmann et al. 2006), we first need to embed it into a population dynamical model with population regulation and a mode of genetic inheritance, and in an “external” environment ***E***. This allows calculation or simulation of “resident” stationary populations where all individuals have the same genotype and strategy ***x***, and following the dynamics of rare strategies with a different genotype and phenotype **x’**. For the sake of easiness of presentation, we assume that there is a single such attractor of the population dynamics Att(***x, E***). Using this model and an Att(***x, E***), we can determine invasion fitnesses of rare mutants **x’**. We assume that a mutation occurs at a single locus and that it has phenotype ***x***’ in heterozygotes. When a mutant allele is rare, the frequency of mutant homozygotes is negligible. We can then write invasion fitnesses as functions s(***x***’, Att(***x, E***), ***E***). For a fixed ***x***, we can call an environment ***E*** an evolutionarily singular environment ESE when

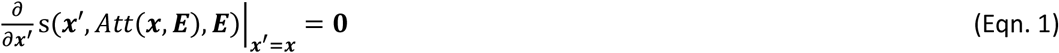

This is a vector of partial fitness derivatives towards each of the phenotypic traits in mutants, evaluated at the resident trait values ***x***. When this vector is zero, no mutant with phenotype ***x***’ slightly differing from ***x*** in one or several traits can invade.

This definition does not suggest that ***E*** evolves. ***E*** does not have to be the environment in which an organism supposedly became adapted. It is just singular. If multiple attractors of the population dynamics occur for an ***x***, the definition needs to be checked in each of them. Adaptive dynamics (Metz et al. 1996, Geritz et al. 1998) usually assumes that ***x*** evolves in a given ***E***. Trait vector **x**^**\***^ is called an evolutionary singularity or evolutionarily singular strategy ESS, when Eqn.1 holds for it, for a given ***E***. Eqn (1) is a first condition for ***E*** to be an environment where ***x*** is adapted and is expected to stay adapted. It can then be verified using adaptive dynamics in a given ***E*** whether resident strategies which are slightly different from ***x*** would be replaced by strategies more similar to it and whether evolution halts when ***x*** is the or a common strategy in the population (Geritz et al. 1998).

### The environment

The temporary pond is assumed to be a disc with a fixed center. The maximum radius which can be covered by water is *r*_max_ (Table 1). The state of a pond is characterized by how much of its extent is covered by water. All non-negative radiuses within [0, *r*_max_] are allowed. The radius covered by water *r*_pond_(*t*) can change gradually and deterministically per day, where *t* can be any date. When *r*_pond_(*t*) = 0, it is in the dry state. In a wet state with positive *r*_pond_(*t*), water is present and fish can hatch, grow and reproduce. When they reproduce, adults lay their eggs in the part of the pond soil which is wet. Depending on the location of an egg within the pond extent, dry conditions start at different times. Location is therefore a state variable of each individual egg. Location states were rounded down to arrive at a limited number of discrete location classes to model (Supplementary Material).

**Table 1.**
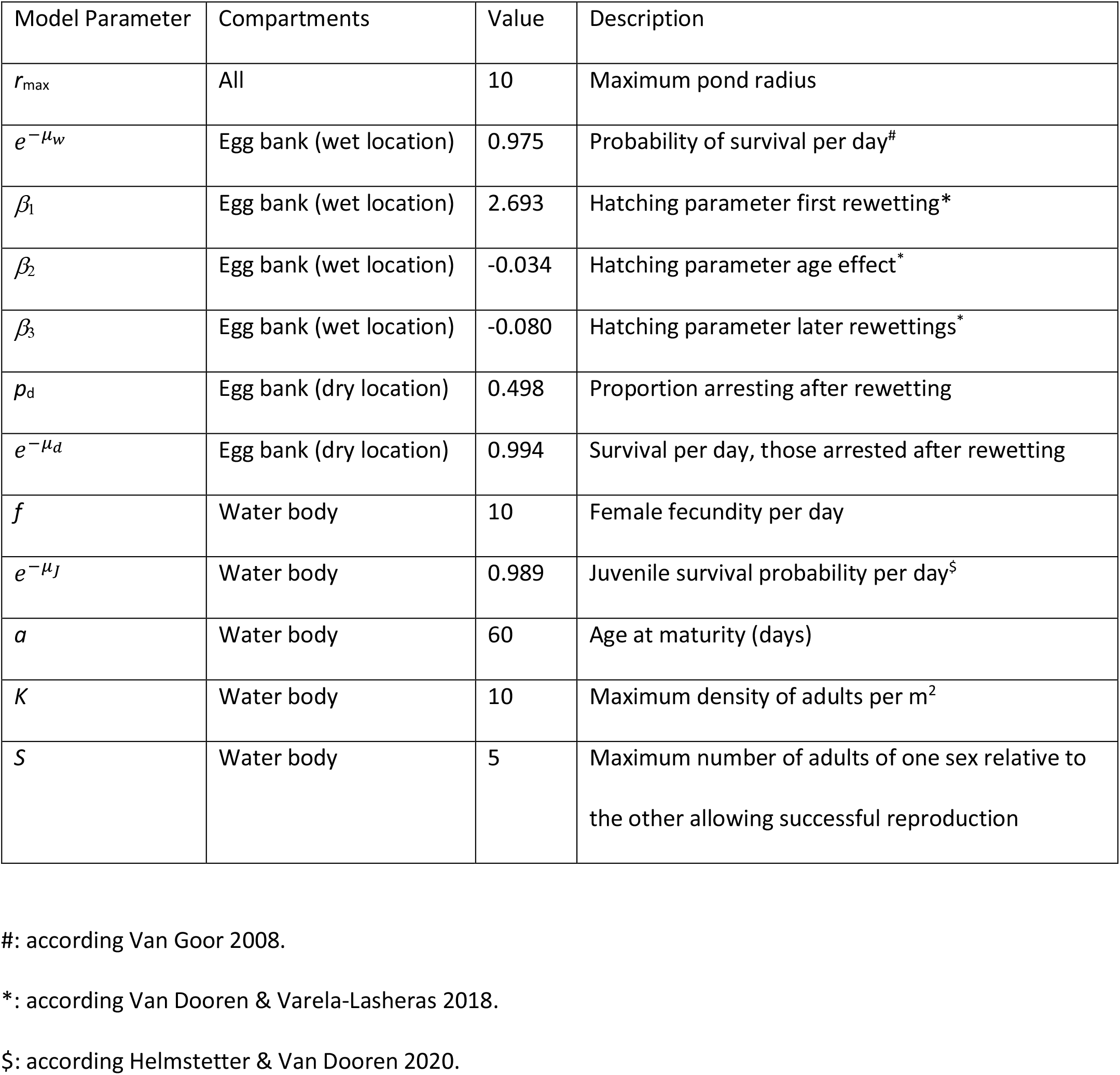
Model parameters fixed across simulations.

| Model Parameter | Compartments | Value | Description |
| --- | --- | --- | --- |
| $r_{\max}$ | All | 10 | Maximum pond radius |
| $e^{-\mu_w}$ | Egg bank (wet location) | 0.975 | Probability of survival per day<sup>#</sup> |
| $\beta_1$ | Egg bank (wet location) | 2.693 | Hatching parameter first rewetting<sup>*</sup> |
| $\beta_2$ | Egg bank (wet location) | -0.034 | Hatching parameter age effect<sup>*</sup> |
| $\beta_3$ | Egg bank (wet location) | -0.080 | Hatching parameter later rewettings<sup>*</sup> |
| $p_d$ | Egg bank (dry location) | 0.498 | Proportion arresting after rewetting |
| $e^{-\mu_d}$ | Egg bank (dry location) | 0.994 | Survival per day, those arrested after rewetting |
| $f$ | Water body | 10 | Female fecundity per day |
| $e^{-\mu_J}$ | Water body | 0.989 | Juvenile survival probability per day<sup>§</sup> |
| $a$ | Water body | 60 | Age at maturity (days) |
| $K$ | Water body | 10 | Maximum density of adults per m<sup>2</sup> |
| $S$ | Water body | 5 | Maximum number of adults of one sex relative to the other allowing successful reproduction |

<sup>#</sup>: according Van Goor 2008.
<sup>\*</sup>: according Van Dooren & Varela-Lasheras 2018.
<sup>§</sup>: according Helmstetter & Van Dooren 2020.

Pond radius profiles over time *r*_pond_(*t*) were kept relatively simple, with up to two wet periods within a year and with linear radius increases and decreases within a wet period. In longer simulations, patterns within a year were repeated across years. In leap years, *r*_pond_(*t*) of the previous year was copied into January 1 to December 30 and December 31 of the previous year was duplicated. To gain a first insight in the effects of pond regime on population dynamics, a small number of seasonal scenarios were explored with a single wet period allowing reproduction (see Figure two for examples). In part of these scenarios, this period was preceded by a wet period too short to allow reproduction. In the simplest scenario (Y0), the pond had water starting on April 11 (day 101), increasing to maximum size *r*_max_ on July 19 (day 200) and became dry again on October 28 (day 301). In scenario Y0, the pond covers 0.3*r*_max_ on day 130 and 0.7*r*_max_ on day 170. From this scenario the period from April 11 to July 19 was then modified, making the period where most of the pond extent is covered by water smaller or larger. Deceleration scenarios were constructed where there was an initial phase of fast filling where 0.7*r*_max_ would be reached on days 163 (YD1), 150 (YD2) and 145 (YD3), respectively. This was followed by a second phase of slow filling with a time span of 37 (YD1), 50 (YD2) and 55 (YD3) days respectively to reach the maximum size *r*_max_ by 19 July. Scenarios with accelerated filling were constructed where 0.3*r*_max_ would be reached after a period of slow filling on days 136 (YA1), 148 (YA2) and 163 (YA3), followed by faster filling, with *r*_max_ still reached on July 19. In a second category of modifications of Y0, a short wet period was inserted from days 71 to 90, with the maximum pond radius within that short period reached on day 80 at values *mr*_max_. Scaling parameter *m* varied from 0.2 to 1 in steps of 0.2, yielding scenarios YS1-YS5. Refilling of a partly desiccated pond was not considered for now and neither was between-year variability. Other environmental state variables than *r*_pond_(*t*) were not modelled. The model does not contain explicit temperature effects yet. Diurnal variation is prevalent in annual fish habitats (Žák & Reichard 2020) but not explicitly modelled here.

**Figure two.**
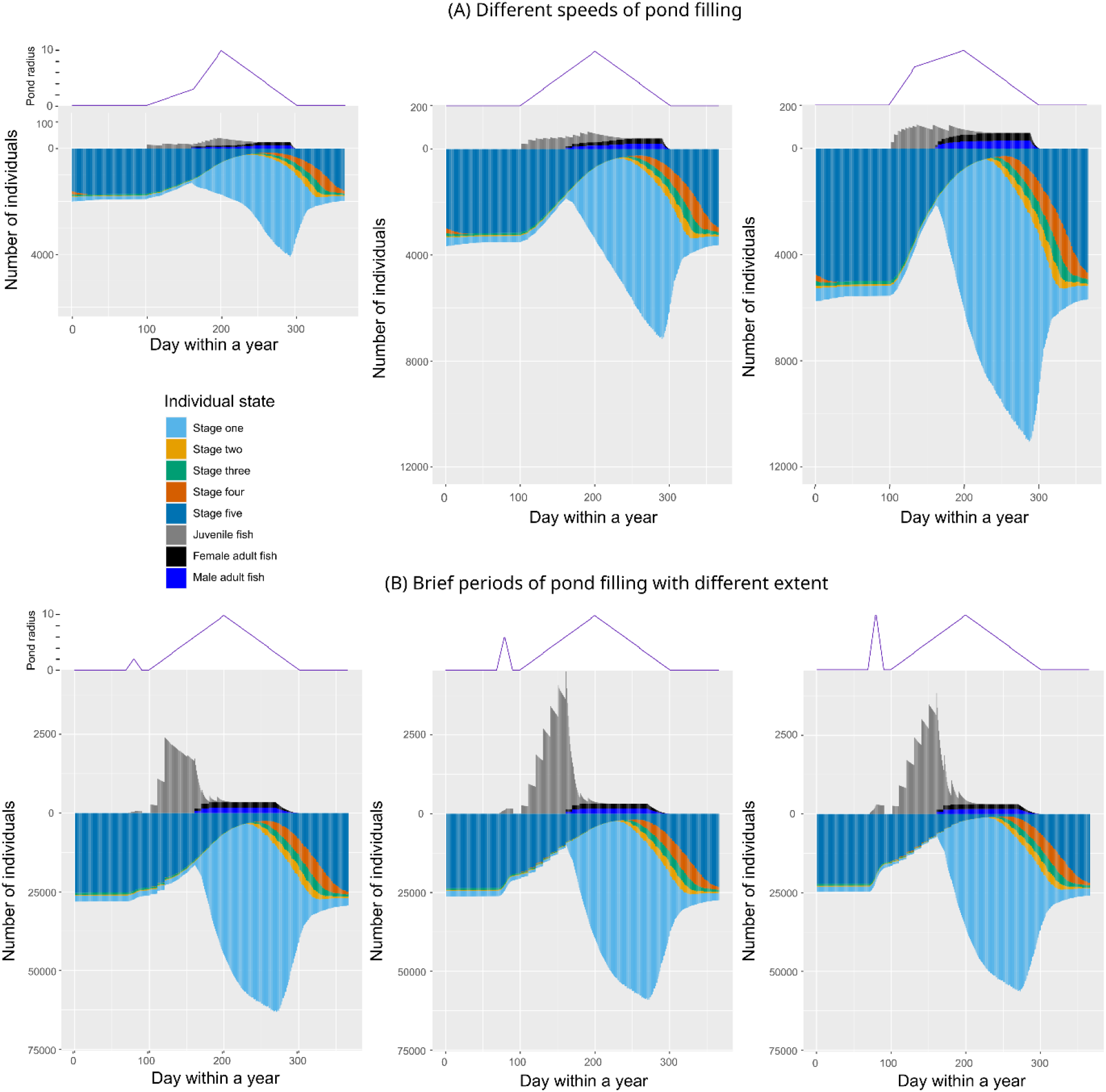
Population dynamics of embryos and fish from day to day in different pond filling regimes. The environmental pond filling regimes are shown above each panel, with the pond radius varying between zero and *r*_max_ = 10. The population dynamics was first iterated for twenty years, then it was checked that it had become stationary. The stationary result was iterated from day to day over one year and the numbers of individuals in different states are plotted. The numbers of individual embryos in the egg bank are plotted (increasing downward) and fish numbers (increasing upward). Note that fish and embryos are drawn with different scales. The bars are as high for one fish as for ten embryos. (A) Effects of changes in the speed of pond filling within a wet period. From left to right, the scenario with accelerating filling YA3, the reference scenario Y0 and the scenario with decelerating filling YD3. (B) Effects of the main wet period where reproduction can occur being preceded by a short wet period which does not allow reproduction. From left to right, scenarios YS1, YS3 and YS5 where this short period covers *m* = 0.2, 0.6, and the full extent of the pond. Note the additional differences in scale between rows (A) and (B).

### Fish life histories

Fish are assumed to move freely within the available water body. Parameter estimates are based on Helmstetter & Van Dooren (2019) and Van Dooren & Varela-Lasheras (2018). Individual fish have their age since hatching and sex as state variables. The number of fish per state and their produced eggs are modelled on a day-to-day basis. Different events which can happen on the same day are treated sequentially. The sex ratio among the hatchlings on each day is equal. Juveniles mature at age since hatching *α*. A simple model of population regulation is adopted. The number of fish within a pond is limited by its surface. It is assumed that each square meter of wet surface can contain a maximum of *K* adults. When more than the maximum number of allowed adults is present on a day (*K* times the area of the pond), surplus adults are chosen at random and erased from the population state. There is no other source of daily mortality among adults during a wet period, senescence is not expected for the wet season lengths modelled (Liu and Walford 1990). When the number of adults *N*(t) is below the maximum which a pond allows, juveniles can survive with a survival probability per day equal to the proportion of surface which is not “occupied” on that day [1 – *N*(t) / (*π K r*(t)^2^)]_≥0_ times a density-independent survival probability 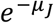. Reproduction occurs each day after density-dependence. Adult females have a fixed individual fecundity per day of eggs *f* which are all fertilized when there are males present in the pond on that day after competition and the sex ratio (ratio numbers of females/males) is within the interval [*S*^-1^, *S*]. Otherwise, fecundity is set to zero because of presumed sperm limitation in males or male aggression. Eggs are deposited at a random location within the pond (Supplementary Material). At the start of a dry period, the remaining adults die. Parameter estimates *α, f, μ*_*J*_ (Table 1) are based on Helmstetter & Van Dooren (2019) and Van Dooren & Varela-Lasheras (2018).

### Simulations of population dynamics

As the extent of the wet pond area changes continually, numbers of individuals are projected forward, not their densities per unit of area (Supplementary Material). Given an initial population state, an egg bank with a number of embryos or a fish population can be projected forward over a series of successive years, where each year repeats the pond cycle of the previous year, to find the attractors of the population dynamics. On these attractors, population states change within a year, but on Poincaré sections at a specific day within the year, the population states can have fixed-point or non-equilibrium attractors. Only fixed-point attractors were observed.

The model for the pond environment, population dynamics and invasion fitness is coded in R (R Core Team 2020) and the scripts are available on https://doi.org/10.5281/zenodo.10992414.

For the purpose of illustration, forward projection was done for example profiles Y0, YA, YD and YS and the stationary dynamics over one year was then simulated on a day-to-day basis.

### Mutational effects

To implement the check of adaptation proposed by Eqn. 1, we need a vector of phenotypic trait parameters ***x*** and the effects of mutations. The embryonic annual fish strategy considered consists of four stage-specific developmental transition parameters and two hatching parameters. The rates are all age-dependent strategies, which would make them four function-valued strategies to assess (Kirkpatrick & Heckman 1989). I opted for a simplification. Four stage-specific trait parameters represent multiplicative changes to the cumulative integrated stage-specific rates (Supplementary Material). The trait values are all one for resident individuals and can be smaller or larger than that for mutants, which then develop slower or faster out of the stages where their trait value differs from value one. For hatching, multiplicative changes to hatching parameters *β*_1_ and *β* _3_ of the linear predictor in the logistic regression are the traits. When the trait values are all fixed at value one, the transition probabilities and hatching probabilities determined from the data are unchanged and represent the assumed common strategy in a resident population of this species. The Supplementary Material details how modified rates affect the transition probabilities between stages.

### Invasion analysis

It is assumed that there is no genetic variation in the resident population with phenotypic effects. For a mutation which is relatively rare, the number of mutant alleles equals the number of mutant individuals because these are approximately all heterozygotes at a single locus of a mutant and a resident allele. Offspring numbers are determined by female fecundity and the availability of adult males. Mutant adult females contribute the mutant allele to half of their eggs. Male adult mutant individuals contribute their mutant allele to half of the eggs they fertilize each day, which is assumed to equal the total number of eggs produced on that day (*f* times the number of resident adult females present), divided by the resident males present.

To estimate invasion fitness of a specific mutation with phenotype ***x***’, the population dynamics of the data-based resident strategy ***x*** was simulated to obtain so-called resident stationary regimes (Supplementary Material). Then the mutant genotype was introduced and simulated competing against the resident strategy over eight complete dry-wet period cycles. The final number of mutant individuals was divided by the initial number at the start of this time interval and raised to the power one over the interval length (in days) and the result log transformed. Invasion fitness is thus the log geometric average population growth rate per day of this mutant population given an environmental regime and a resident data-based population strategy.

### Fitness sensitivities

To assess the effects of mutants with specific trait changes on invasion fitness in a given environmental regime, mutants with trait values between 0.9 and 1.1 were generated for each of the separate traits and their invasion fitnesses were estimated in an established population with the data-based strategy. This was done for each of the pond filling scenarios Y0, YA, YD and YS. Per environmental regime, the invasion fitnesses for mutations in each trait with value 1.1 were used as a measure of invasion fitness sensitivity (Caswell 2019) and plotted to see how environmental properties might affect strengths of selection and change the potential adaptedness of strategies.

For the five stage-specific mortalities, invasion fitnesses were also calculated using the same approach as for rates of development. This allowed a comparison of strengths of selection between developmental and mortality rates. To compare these strengths of selection more easily between environments, we assigned parameter values to the scenarios Y0, YD and YA, and to Y0 and YS, such that invasion fitnesses of a specific trait change could be plotted as functions of environmental parameters. YD1-3 were given parameter values -1, -2 and -3, Y0 value zero and YA1-3 the values 1, 2 and 3. To compare effects of adding a short wet period where reproduction is impossible, Y0 was again assigned parameter value zero, and YS1-5 the corresponding values of *m*.

### Evolutionarily Singular Environments

A more detailed parametric specification of pond filling regimes (Supplementary Figure one) was investigated to search for ESE. Here, a parameter vector *ρ* contained ten parameters *ρ*_i_ with values between zero and one, affecting dry and wet interval lengths and maximum pond radiuses within intervals. A search algorithm on vectors *ρ* was implemented to investigate whether the data-based strategy might be adapted to one or some of these. The Euclidian norm of the vector of invasion fitnesses of mutants with trait values 1.1 in the four developmental traits and in the two hatching probabilities was calculated for each environmental regime visited by the algorithm as a measure of the selection gradient. These gradients were minimized as a function of *ρ* by means of the L-BFGS-B method (Byrd et al. 1995), programmed in the optim function for R (R Core team 2020). In an ESE environment *ρ*^*^ where the data-based strategy is adapted, this norm would become zero.

Optimizations were started from different initial pond regimes with one or two wet periods, also with two wet periods long enough for reproduction to occur. In all cases explored, the algorithm converged to a local minimum of the gradient of invasion fitness. Its initial and final values at these local minima were calculated. Environmental regimes with the smallest gradients found are plotted, their resident population dynamics simulated and shown, as well as the pattern of invasion fitnesses of mutants in these environments.

## Results

### Population Dynamics

Pond regime differences within years can change the expected number of individuals in the egg bank more than tenfold (Figure two). Fish are always a minority among the individuals present in the population (Fig. 2). Different speeds of pond filling make periods with many reproducing adults shorter or longer. Coexistence of adults and juveniles can occur over extended periods (Fig. 2). Their numbers vary with pond extent, but the number of adults is not always matching the profile changes closely. It seems that the strong competition effects of adults on juveniles prevent this. During dry periods all embryonic developmental stages are usually present but stage five is generally most abundant (Fig. 2). At different distances from the pond center, the number and distribution over stages of embryos change predictably, with more advanced stages found sooner at larger distances from the pond center (Supplementary Figure two). At the start of a dry period after reproduction, earlier stages are the most abundant near the pond center and more advanced developmental stages dominate near the pond edge (Supplementary Figure two).

The largest egg bank changes occur within wet periods, due to hatching, overall lower survival, and the addition of new eggs. Variable speeds of pond filling have clear effects. With faster initial and slower later filling as the maximum radius is approached, the overall size of the egg bank increases at all times because more individuals reproduce (Fig. 2A). For comparison, in pond regimes with step functions between no water and a full pond, populations go extinct when the wet season is shorter than 189 days (results not shown). The increase in number of embryos accelerates or decelerates as the pond radius does (Fig. 2A), but there cannot be a causal relationship, as a deceleration also occurs for other pond regimes where a short wet period precedes the one with reproduction (Fig. 2B). Accelerating egg number changes with increasing pond size might be indicative of a lack of competition between mature fish. With an additional short pre-period of pond filling covering the pond radius to some extent (Fig. 2B), stage five embryos hatch in that period without any chance of reproducing and the embryos which experienced their first rewetting have their hatching state variable changed. In the main wet periods (Fig. 2B), we can see more intense competition: the fraction of the juvenile population contributing to the pool of reproducing adults is smaller, while overall, there are many more new eggs produced. If we need to guess which among these environments might be the most adapted, based on simple principles such as the optimization of population size (Charlesworth 1994) or the pessimization of the environment (Metz et al. 2008), it would be the pond filling regime where the egg bank is largest and where competition is most intense, so YS1 or YS2. A combination of decelerated pond filling and a short wet period before a long one might be the environment where the data based strategy is the most adapted, if only within year variation would occur.

### Invasion fitness patterns

When invasion fitnesses are estimated for changes in each trait in each of the example scenarios, it can be observed that invasion fitness changes are largest for mortality and development in stage one, and for early hatching (Figure three). Note that for mortalities in other stages than stage one, the slope of invasion fitness on mutational trait change is small. This can be explained by the already high survival rates in each of these stages. For the development rates, there seems only weak selection to increase developmental speed in most stages (Fig. 3). However, selection was always towards acceleration. Invasion fitness slopes changed when environmental parameters changed, in particular for the traits under stronger selection (Figure four). For the range of changes explored in (Fig. 4), invasion fitness slopes did not change sign except for hatching at the first rewetting, when short wet periods without reproduction were added to the reference scenario Y0. In these YS pond filling regimes, parameter *m* was varied gradually to find an ESE but it was not present, at value *m* = 0.1, selection changed direction abruptly.

**Figure three.**
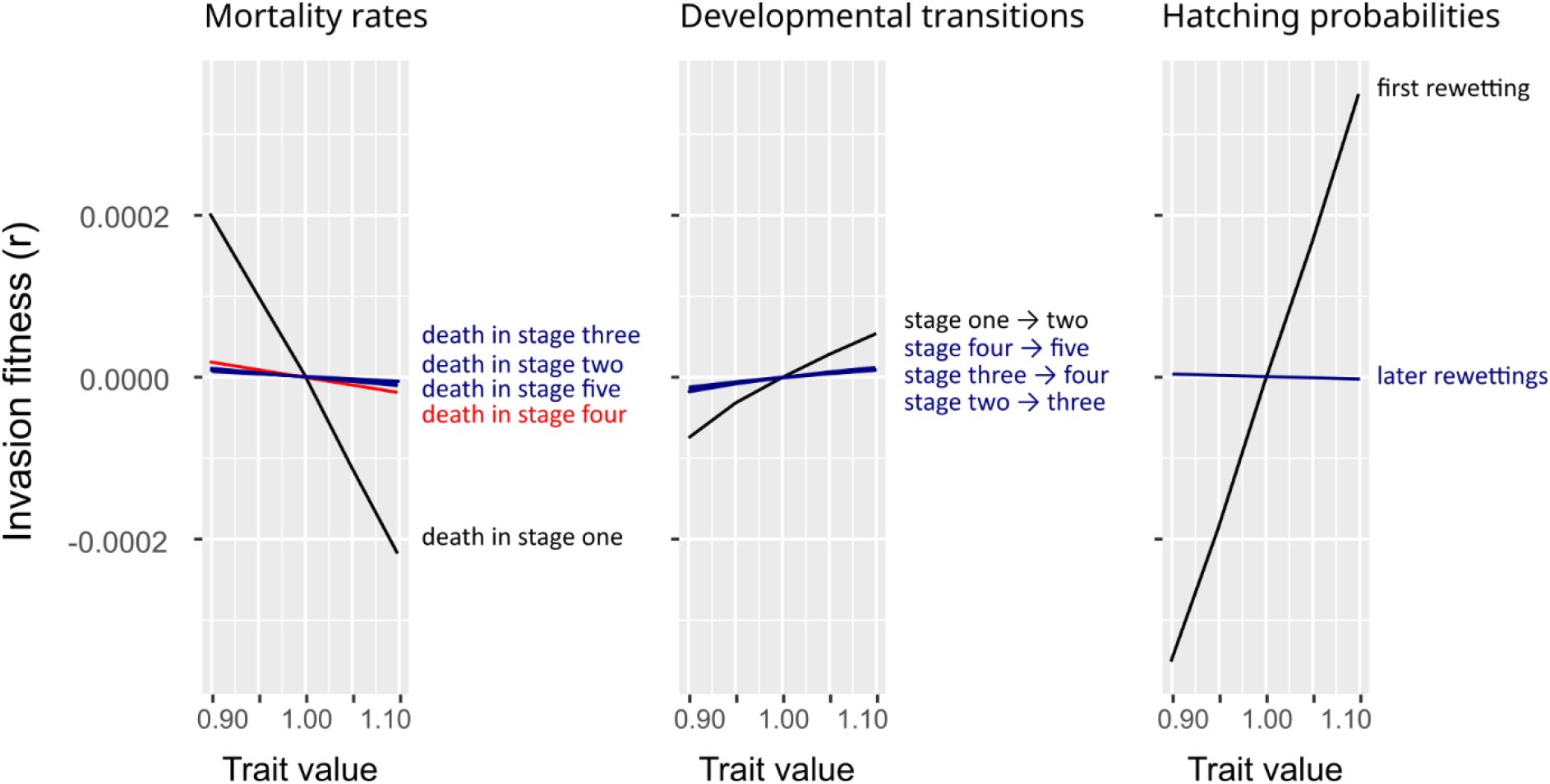
Invasion fitnesses of phenotypic mutational changes in each of the embryonic life history traits, in the simplest pond filling regime Y0. Invasion fitnesses are shown for invasions of mutant phenotypes in a resident population which has become stationary in scenario Y0. The resident trait value for each trait is at value one. Invasion fitnesses are estimated at five equidistant points and connected by a line.

**Figure four.**
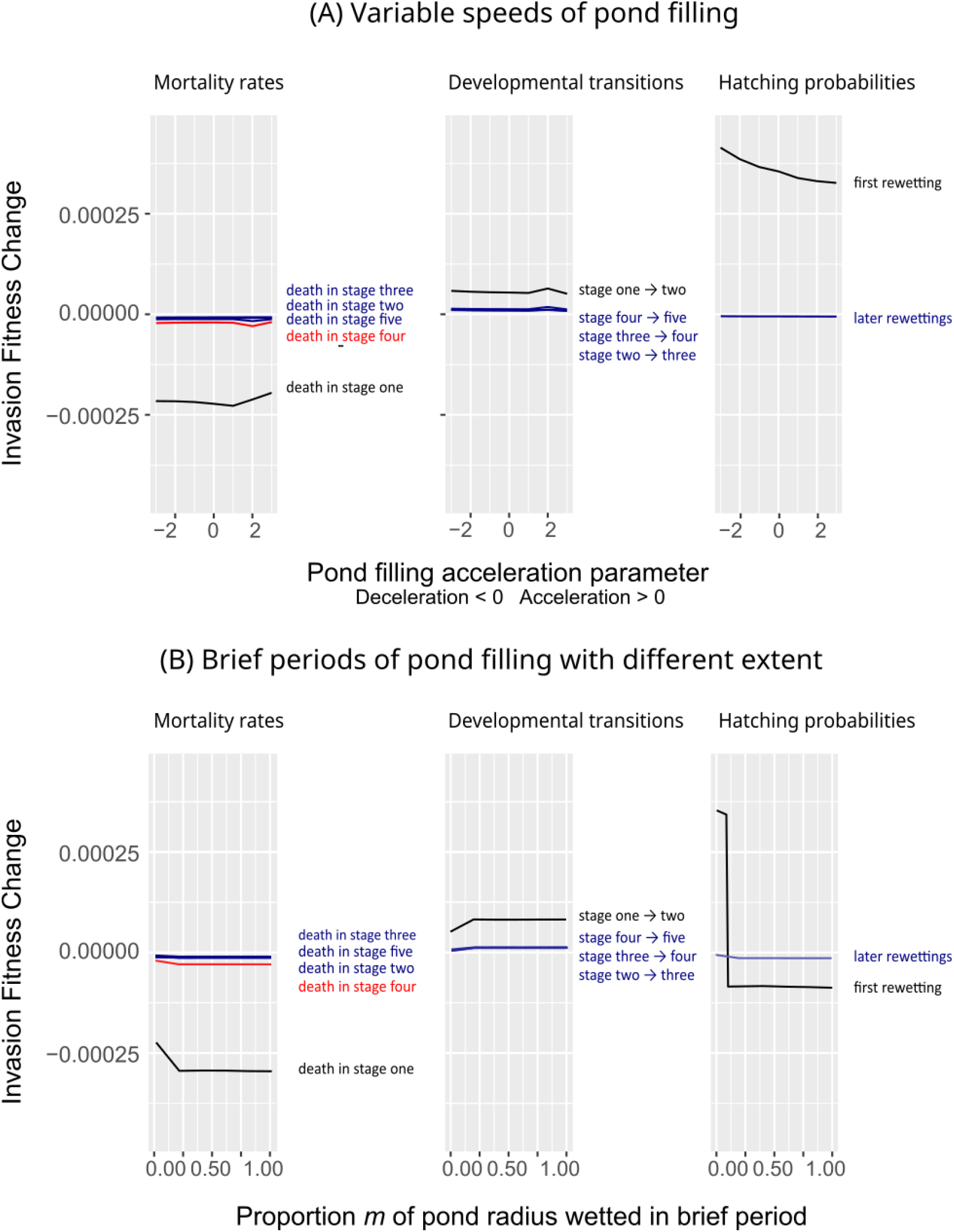
Sensitivities of invasion fitness to pond regime changes seem limited for many traits. Patterns of invasion fitness for a fixed difference between mutant phenotype and resident phenotype are shown. Invasion fitnesses of mutants increasing cumulative hazards and logistic regression parameters with a multiplication factor of 1.1 plotted as a function of an environmental parameter. This gives a qualitative impression of strengths of selection in different environmental regimes on each of the traits. Labels of rates next to panels are ranked according values. (A) Decelerating and accelerating pond filling. At zero on the x-axis, the pond filling regime is the one of the middle panel in row A of Figure 2. (B) Short wet periods with increasing fractions *m* of the egg bank covered. For hatching at first rewetting, it was determined in a stepwise manner for which value of *m* the sign of the sensitivity changed. The change was abrupt.

### Evolutionarily Singular Environments

It was found that the minimization algorithm could make jumps to environmental regimes where dry periods would disappear and the population went extinct. These iterations were restarted with a constraint which imposed that a dry period would exist for of at least 10% of the year. In general, initial conditions with two periods where reproduction could occur were replaced by pond regimes with a single wet period with reproduction. Therefore, the annual fish strategy here is not adapted to regular pond regimes with two wet periods within a year where reproduction occurs. When pond regimes were changed iteratively such that the invasion fitness gradient for the developmental speeds and hatching traits became minimized, the output pond regimes to which the algorithm converged were local minima which had filling regimes with a subset of parameters modified. Figure 5 shows two representative initial and final pond filling regimes. In general, the length of the invasion fitness gradient vector would become at least halved, but vectors of zero length were not obtained. Therefore, we did not find evolutionarily singular environments with within-year changes and variation only. The only trait which ever became really adapted in one of the environments, was late hatching (Fig. 5C, lower panel). Minimization of the invasion fitness sensitivity in early hatching, a trait which contributed much to the reduction of the gradient, could co-occur with an increase in invasion fitness sensitivities for other developmental speeds (Fig. 5C). The attractors of the population dynamics in each of the regimes at local minima of the invasion fitness gradient, had large maximum numbers of eggs in the egg bank and showed that many juveniles did not become reproducing adults. Note that the output example in the first row has a larger number of eggs in the egg bank than any of the other pond filling regimes investigated. At the start of the first wet period in the year, the two scenarios differed in number of eggs in the egg bank by a factor of six. In all, these simulations also found that faster developmental transitions were always favoured which replicates the standard conclusion for “constant” environments.

**Figure five.**
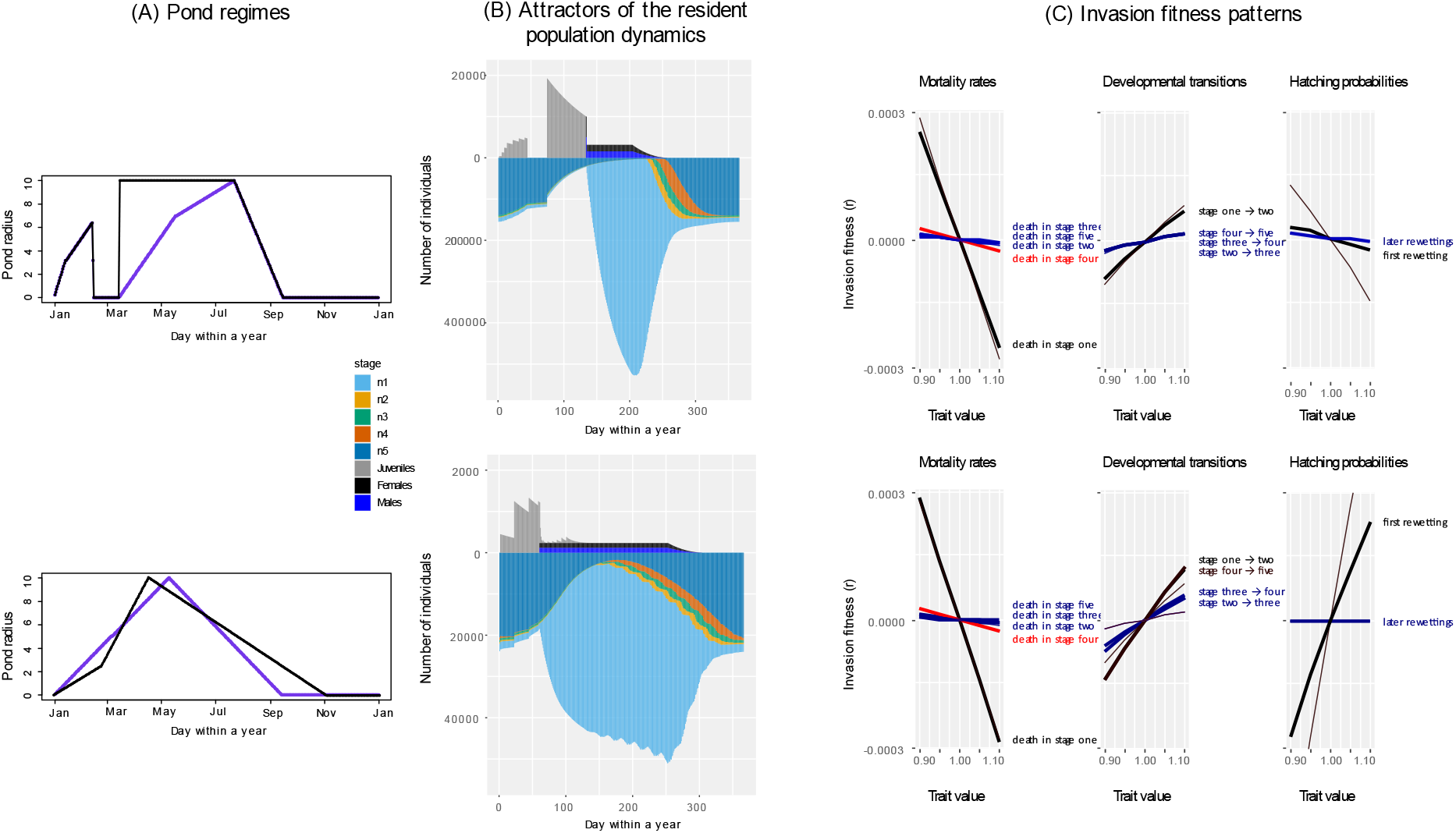
Minimization of invasion fitness gradients by changing pond filling regimes results in limited reductions of selection strengths. Results of optimizing environmental pond filling regimes such that the data-based strategy becomes as adapted as possible, by a parameter search. (A) Two initial pond filling regimes from which optimization started (purple), the upper one with two wet periods, the lower profile with a single one. Resulting pond filling regimes after minimization of the invasion fitness gradient are shown in black. (B) Attractors of the resident population dynamics in each of the two pond regimes to which the algorithm converged. Note that fish and embryos are drawn with different scales. The bars are as high for one fish as for ten embryos. Both rows additionally have different scales. The population sizes are much larger for the first row. (C) The patterns of invasion fitnesses per trait in the regimes obtained after minimization. The invasion fitnesses in the initial pond regimes are shown as hairlines for comparison. Traits with very similar invasion fitness patterns are given the same colour. Labels of rates next to panels are ranked according their values. It can be seen that for some traits, the invasion fitness slope increased while the multi-trait fitness gradient became minimized.

For the output pond filling regime in the second row in Fig. 5, the evolutionary stability of hatching at later rewettings *β*_3_ was investigated and it was found to be an evolutionary repeller (Geritz et al. 1998). For the output example in the first row in Fig. 5, pond filling regimes were further optimized as a function of the invasion fitness sensitivity on the hatching at first rewetting trait *β*_1_ alone. A pond filling regime was found (with *ρ*_8_ = 0.68) where that sensitivity vanished, but it was found that the parameter value in the data corresponded to an evolutionary repellor (Geritz et al. 1998).

Therefore, even while ESE can occur for observed hatching probabilities, these have no long-term evolutionary stability in these environments. For values of *m* above 0.1 in the YS pond filling scenarios, an exploratory analysis was carried out. It was investigated to which evolutionarily singular value evolution in *β*_1_ would converge, which turned out to be values which corresponded to very low probabilities of hatching at first rewetting. When this was done for *β*_3_ it could either converge to higher or lower values, indicating for each *m* that there was an evolutionary repeller nearby the data value for this trait. There is therefore no evidence that the intermediate hatching probabilities are adapted to within-year pond filling variation, this analysis suggests that all-or-none hatching probabilities would be adapted.

## Discussion

For an embryonic life history strategy of annual fish (Van Dooren & Varela-Lasheras 2018), it is investigated to which environments it might be adapted. For this, the observed strategy is complemented with a model for the environment and the annual fish population dynamics in it. This allows searching for Evolutionarily Singular Environments ESE where the derivatives of invasion fitness with respect to the strategy trait parameters are all zero at the observed strategy. This is a first condition for being an evolutionarily stable strategy ESS in a given environment, and can therefore be used to single out environments to which the observed strategy could be adapted. The analysis did not find an ESE among the regimes considered in which all six traits would be adapted. Only for the two hatching probabilities, ESE pond filling regimes without any between-year variability were found. However, the hatching probabilities did not have evolutionary stability there and might remain some time unchanged by selection, but not so I the long term. Given model assumptions, an environment with regular within year variation and one or two periods where reproduction can occur does not seem to be the environment to which the embryonic rates of development and hatching probabilities of these annual fish are adapted.

### Adaptation to deterministic pond size variation within years

In a sense, within-year stochastic variability in demographic parameters is present. Random egg deposition within a pond combined with the arrival of a wet period too short for reproduction reduce the correlation between individual fitnesses of randomly chosen pairs of individuals (Frank & Slatkin 1990, Starrfelt & Kokko 2012) but also increases their variance in reproductive success.

Hatching probabilities could be bet-hedging traits adapted to this within-year uncertainty, Selection for smaller hatching probabilities at first rewetting than the observed one was advantageous in cases where the wet period where reproduction could occur was preceded by a shorter period during which juveniles could not mature. In one such case, by changing the extent of the pond which was covered by water during this period, an environmental regime was found which minimized the effect of this hatching probability on invasion fitness. In this ESE and a different one for hatching at later rewettings, the hatching probability traits were evolutionarily unstable. The suggestion by Pinceel et al. (2015) that observed hatching probabilities would be adapted to short-term environmental uncertainty could not be confirmed for this system. An exploratory analysis rather suggested that the evolutionary dynamics in a given environment would converge to a combination of very low and high hatching probabilities and therefore no diversified bet-hedging.

In the study of adaptation to environmental variability, most attention has been given to uncertain environments (Tuljapurkar 1990) and not to deterministic patterns of change as modelled here. The reference environment contrasted with uncertain environments is usually called “constant” (Stearns 2000, Rose et al. 2002). If this refers to demographic parameters being invariable across years, deterministic seasonal patterns can be implicitly present and even randomness at the individual level. The boundaries of seasons often define the moment of census (Venable 2007). It would be better to focus on models where the underlying demographic processes and the observation process can be uncoupled, such that we can predict population states at any census point, to be chosen freely within a year and keeping seasons explicit (e.g., Cooch et al. 2003). Phenological changes which shift seasonal boundaries now occur often due to climate change (Park & Post 2022) and this then generates between-year variability (ten Brink et al. 2020). This can occur without differences in instantaneous rates, just by duration variation.

Not all possible within-year deterministic changes were explored, such as refilling of a pond after a partial desiccation. This is postponed until data on competition between juveniles and adults and on survival of different stages in wet conditions become available, which both are expected to affect patterns of selection. When periods where a partially desiccated pond refills occur, different stage-specific mortality rates in wet hypoxic conditions might switch the direction of selection on development such that stages with increased mortality in wet hypoxic conditions are avoided. A state switch such as between advancing/non-advancing states might be of use then. Note that low rates of development in stages two and three and diapause II, claimed to be the hallmark of annualism (Furness et al. 2015a, Furness 2016), were not observed in the data (Van Dooren & Varela-Lasheras 2018). Assuming that this absence is adaptive, the presence of diapause I suggests that mortality in wet hypoxic conditions might increase for this species in stages two or three where diapause II occurs, rather than decrease (Podrabsky et al. 2007).

In some of the simulation steps aiming to minimize the invasion fitness gradient, pond regime vectors *ρ* were chosen without a completely dry period. These resulted in population extinctions, but might be useful to explore the adaptedness of developmental patterns in non-annual killifish (Varela-Lasheras & Van Dooren 2014) and why killifish clades which evolved annualism never seem to have lost it. On the other hand, regimes with two periods where reproduction could occur always had larger fitness gradients than regimes with a single such period, such that no support for adaptation to a second chance to hatch and reproduce within a year (Furness et al. 2015, Polačik et al. 2017) was found.

### Adaptation to variation between years

In models for annual plants, dormancy hedging seed germination over years could substitute for a within-year bet-hedging trait (ten Brink et al. 2020). Therefore, bet-hedging in hatching probabilities might still not be adaptive when years differ. A common conclusion has been that risk avoidance by bet-hedging in one trait, would pre-empt the scope for bet-hedging in other traits (e.g. Koons et al. 2008, ten Brink et al. 2020). This is however, only partly supported (Snyder 2006). Many of these conclusions derive from models where individuals could hatch or germinate any time within a year or every year (Metcalf et al. 2015, Poethke et al. 2016, ten Brink et al. 2020), which is different from how annual fish interact with their environment. In annual fish, hatching is provoked by the arrival of water. In dry years, hatching is impossible. If long-term survival in the pre-hatching stage is reduced relative to earlier stages, individuals should become spread out over years earlier in development and stage-specific survival would constrain hatching to spreading individuals over within-year occasions only.

Delaying and bet-hedging can be selected against in any constant environment or with recurring within-year deterministic environmental changes, but become advantageous with different population fluctuations. This could be stochastic variation, when random samples of annual pond regimes are concatenated, or rather deterministic time series with population size fluctuations (Bulmer 1984, Van Dooren & Metz 1998). Between-year variation in pond filling patterns would only render delaying adaptive if it manages to dampen fluctuations in the densities of embryos in the egg bank.

Evolutionary ecology has a long tradition of studying adaptation to variable uncertain environments (Cohen 1966, Ellner 1997, Stearns 2000, Evans & Dennehy, 2005), i.e., environments with unpredictable random changes. Stochastic environmental variability in pond filling regimes between years will render changes in numbers between years random, even without stochastic variation in instantaneous embryonic life history rates in wet and dry soil. The body of theoretical results on life in uncertain environments then becomes applicable to this system (Tuljapurkar 1990). As this model has explicit links between environment and demography, it would permit relating trait variation to magnitudes of environmental uncertainty (Simons 2011, e.g., Pinceel et al. 2017) or to variances in life history statistics or demographic parameters such as survival or fecundity (Simons 2011, e.g., Venable 2007). In modelling, however, the environmental variation itself has been kept relatively implicit, and the focus has been on random ‘environmental’ life history variation (Rose et al. 2002, Ellner 1997). Between different within-year pond filling patterns shown, the stationary population size in the egg bank can differ by a factor hundred (Figs. 2 and 4) such that there seems scope for pond filling regimes which vary between years to generate egg number fluctuations and for adaptation to dampen these via the adjustment of embryonic life history rates.

Unfortunately, with the current trends in environmental change, it is unclear whether we can still expect real populations to be near attractors of their eco-evolutionary dynamics for much of the time. For example, the recently described *Austrolebias wichi* was at that time known from a single pond in a region with large changes in agricultural practise (Alonso et al. 2018), but can be now found in nearby man-made roadside ditches as well. Similar to the search for environments rendering embryonic life histories adapted as implemented here, simulations of anthropogenic disturbances and rearrangements of the natural habitats could explore which ones would render patterns of development adapted and would minimize adaptational lags (Van Dooren 2022) across a wider range of projected climate change patterns. Such managed novel environmental regimes which are ESE could aid in the conservation of the several hundred annual fish species occurring in South-America and Africa, often at risk of extinction and with currently small population sizes.

### Model validation and falsification

Different types of data can be collected to test model predictions and assumptions. First of all, snapshot distributions of embryos in the egg bank across developmental stages as in Polacik et al. (2023) can be compared to model predictions. The model predicts on many days that a single stage will dominate in the egg bank (Fig. 2 and Supp. Fig. 2), therefore, large sample sizes will be required to reject the model on the basis of the relative occurrences of the less abundant stages. Predictions can also be made for transfers of embryos between lab and field (Polacik et al. 2023) and simulations can be used to optimize power of such experiments. Several model assumptions were made based on the lack of more detailed data, each attempting to avoid introducing more than a minimal amount of parameters. Lacunes in the data call for a field embryology with longitudinal studies in time sequences of natural conditions and also for studies on competition between fish. The field studies of Polacik et al. (2021, 2023) collected data on batches of eggs, not individuals. They could not detect any developmental arrest in the field, but did report data which allows an estimate of survival of embryos in wet soil which would imply that one third of them would be able to survive one year in wet soil. I used a lower estimate of survival based on unpublished individual data, from eggs placed in wet soil one day after they were laid. Estimates of survival in wet soil conditions in other stages than the first are needed and a longitudinal follow-up of embryos after their first rewetting. More data are needed on hatching probabilities after the first rewetting, to estimate age and other effects there. Detailed data on pond filling regimes are needed as well, to investigate what the speeds of emptying and filling actually are, how these vary within a year and what the magnitude is of between-year variation.

### Resources, rates and sizes

Searches for an annual pond filling cycle where embryonic life histories would be adapted halted at environmental parameters where the invasion fitness gradient was at a local minimum. There, invasion fitness sensitivities of developmental rates and hatching probabilities were smaller than in the initial environments but never zero for rates of development. It is necessary to consider that trade-offs might nevertheless make life histories adapted: if mortality rates and developmental rates are coupled by trade-offs provoked by allocation of resources, negative and positive invasion fitness changes might balance due to within-year variation alone. Rates within and between stages might be coupled. For example, Van Dooren & Lasheras (2018) found that when parental pairs were compared, low mortality in stage one was associated with faster development from stage 4 into the prehatching stage. Also constraints might render the observed strategy adaptive. It would then be on a boundary of the allowed trait space. In recent years there have been efforts to understand whether and how plasticity and maternal effects evolve given uncertainty (de Jong & Behera 2002, Dey et al. 2016, King & Hadfield 2019, Van Dooren 2022). Decisions on the allocation of resources might be plastic, and the adaptiveness of amounts of plasticity was not investigated here. We know that desiccation plasticity is limited in the annual fish studied (Van Dooren & Varela-Lasheras 2018), such that it might be little predictive of the future. Insight in temperature plasticity is lacking. In African *Nothobranchius* annual fish, Furness et al. (2015b) demonstrated temperature plasticity in the entrance into diapause. However, these authors defined developmental duration as time to hatching or death, such that separate rates of development were not estimated.

Here, competition between juvenile and adult fish was modelled according a very simple density-dependence. It ignored size-structured competition to a large extent (e.g., de Roos et al. 2003). Size effects can extend to the embryonic stages, if egg size and hatchling size correlate. Vrtilek et al. (2020) found no correlation between them in *Austrolebias bellottii*, while in a different species hatchling size depended in a hypo-allometric manner on egg size. Including more size structure where possible would permit investigating coexistence and co-adaptation of annual fish species (Helmstetter et al. 2020).

### Eco-evo-devo modelling

Developmental life histories were embedded into a demography across the entire life cycle, bringing us closer to an eco-evo-devo which is predictive and can start to relate environmental regimes to adaptation. Four of the six traits modelled summarize effects of mutations across all ages on function-valued development rates. This might not correspond to how specific genes would change phenotypes, but at least most possible events in the embryonic life histories are included in the assessment of adaptedness. The multi-state model used for the embryonic life history has cumulative hazards which depends on the number of days in dry conditions, leading to a Markovian model (de Wreede et al. 2010). This might not be the model which integrates developmental mechanisms easily. Van Dooren & Varela-Lasheras (2018) modelled development with age-within stage as timescales, which seems reasonable when a minimum time is required to complete each developmental stage. With such timescales or coexisting other timescales, the only way to arrive at predictions of distributions of individuals over states is forward simulation of all competing processes (de Wreede et al. 2010). This is individual-based modelling and recalls conclusions of Hogeweg and others in the classical school of bioinformatics (Hogeweg 2012), dealing with the information exchange of organisms and their environments and the incompressibility of this process. Modelling the dynamics of pools of physiological resources and state variables explicitly is an alternative approach which others have implemented to keep track of processes with different timescales (DEB modelling, Kooijman 2010), and this should be explored as well. Embryos within an egg have well-circumscribed resource pools available to them. In the end, multiple such efforts, even for the same empirical model, seem required to understand evolutionary constraints on development and to be able to project adaptation and evolution as processes fuelled by mutations forward into the future. Fortunately, for annual fish, rather mechanistic models are available for processes surrounding diapause I (Reig et al. 2017, Montenegro-Rojas et al. 2023). However, an approach which is customary, even in evo-devo, is to separate the processes generating phenotypic variation from selective processes such as mortality (e.g., Salazar-Ciudad & Cano-Fernández 2023).

This should be abandoned. Individual phenotypes are not composed on a conveyor belt moving individuals to a station where their quality is controlled. They can change the speed of the belt, they can fall off, jump off and back on again. At any point in time there are competing risks of advancing their phenotype and of dying which can vary between genotypes.

## Acknowledgements

This study was supported by action Climat-AmSud 21-CLIMAT-06, funded by CNRS France, CONICYT Chile and ANII Uruguay.

## Supplementary Material

### Transition probabilities

Cumulative hazards are measures of the summed probabilities across all observation times that individuals have made a specific transition by a specific time (de Wreede et al. 2011) and are represented using time-dependent matrices ***A***(*a*), here of age *a*, with *a* ranging from zero to the maximum age observed/considered. ***A***(0) equals zero. Each entry *A*_ij_, with *i* denoting the row and *j* the column of each matrix element, represents the cumulative hazard at that time for the transition from state *i* to state *j*. Note that this is different from standard practise in matrix population dynamics models, where columns denote departure states and rows arrival states (Caswell 2000). Six states can occur in an embryonic life history and are indexed as the five developmental stages (*i, j* = 1…5) and the dead state (*i or j* = 6). All sub-diagonal elements *A*_ij_ are zero. In a statistical analysis, non-parametric estimates of matrix function ***A*** can only change at ages where individuals have been observed. If we give observation times *a*_*τ*_ an index *τ*, then the estimated changes in matrices ***A*** at ages *a*_*τ*_ are given by matrices d***A***(*a*_*τ*_). Transition probabilities from age *a*_1_ to *a*_2_ are calculated as (Eqn. 1, de Wreede et al. 2011)

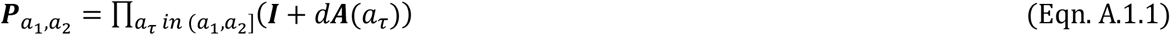

where I use interval limits as in de Wreede et al. (2011). Using the data-based transition probability matrices (Eqn.1), a set of matrices ***P***_*a,a*+1_ is constructed projecting numbers of embryos in different stages and sharing their age state variable one day forward, for any age *a* ∈ [0, ∞]. For all ages above the maximum *a*_*max*_ observed (which was 169 days), it is assumed that ***P*** remains equal to 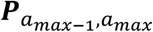. To project numbers of embryos forward in time, we only need to project the numbers of individuals alive. Therefore the 5 by 5 minorized matrices ***M***_66_ are used, with the sixth row and column of ***P*** removed. The life history transitions in this environmental compartment imply that embryos need, next to their location, their age and stage as state variables. However, as it is only age given in the number of days in dry conditions which is assumed to determine developmental progress, we use this number as the individual age variable.

### Population Dynamics

The equations are given for the population dynamics of a single mutant genotype and phenotype across wet and dry periods. The equations include the feedback from the resident population. It is explained how to obtain the population dynamics of individuals with the resident genotype and phenotype using similar equations. In principle, development and mortality are processes occurring or modelled on a continuous timescale, but the periodicity in environmental conditions within a day is strong (Jacobs et al. 2008) and a motivation to use days as the discrete time step to carry out the modelling instead.

Embryos have different individual state variables. Numbers of embryos *N*_ijkl_ are indexed *i* for stage (*i* = 1…5), *j* for their location, i.e., their distance from the pond center in meter (*j* ≥ 0), *k* for the first or later rewetting variable (hatching state *k* = 1, 0) and *l* for the advancing (*l* = 1) and non-advancing states (*l* = 0). Embryos are not differentiated according their sex.

A location *j* is dry when *r*_*pond*_(*t*) < *j* or *r*_*pond*_(*t*) = 0. In dry conditions at their location on day *t*, embryos are projected forward into the next day according

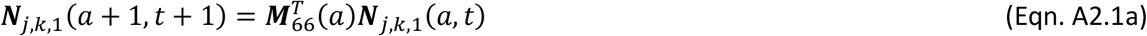

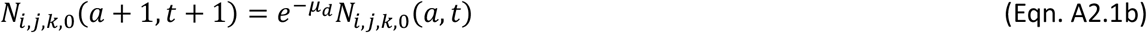

where ***N***_*j,k*,1_ is the vector of numbers *N*_*i,j,k*,1_ of embryos in the five developmental stages. A location *j* is wet when *r*_*pond*_(*t*) ≥ *j*. On the first day of wet conditions at a location *j*, embryos in the pre-hatching stage can hatch. To determine the number of individuals hatching *H*_*j,s*_(0, *t*) at time *t*, with sex *s* and with their age as fish zero, from embryos of all ages and per location *j* receiving water, we need to use the hatching probability functions and account for the fact that half of the individuals are assumed to be females, half males. Index *s* equals *f* for females and *m* for males. When a location is already wet for at least one day *H*_*j,s*_(0, *t*) is zero. For a category *j* which is wetted

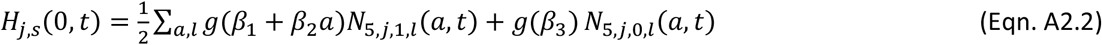

The individuals which did not hatch at *j* are

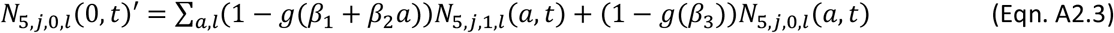

Since these individuals all experienced a rewetting now, the index *k* is set to zero for all of them and *N*_5,*j*,1,*l*_(0, *t*)′ equals zero for both *l*. Then on the same day, the advancing and non-advancing states are reset and the age state variable, the number of days in dry conditions, is reset to zero. For stages one to four this yields

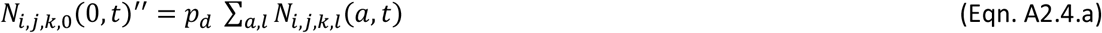

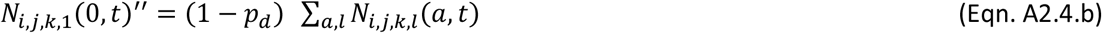

with *N*_*i,j,k,l*_(0, *t*)^′′^ the number after resetting at location *j*. For stage five, this occurs in the individuals remaining in the egg bank after they decided not to hatch

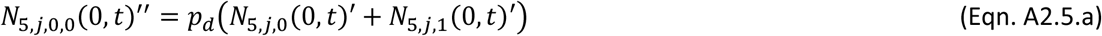

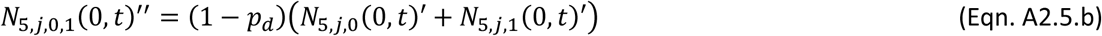

The reset individuals *N’’* are then advancing to the next day according the survival probability in wet conditions

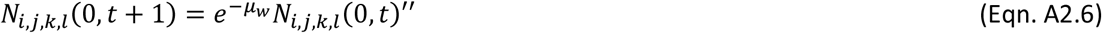

and on subsequent wet days at location *j*, individuals are updated according

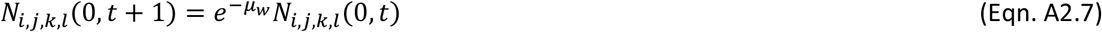

Fish can be present when *r*_*pond*_(*t*) ≥ 0. The fish population is initiated from hatchlings.

The number of juvenile fish of age one are the summed individuals that decided to hatch on the previous day. For older ages (*a* below *α*), they are the juveniles which survived from the previous day, including the effects of density dependence from the resident population

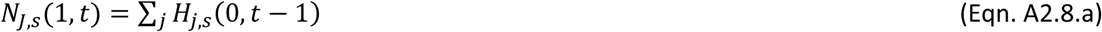

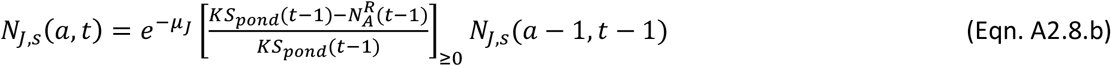

where 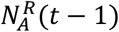 is the number of adult resident individuals present at the start of day *t* – 1 and *S*_*pond*_ (*t* − 1) the pond wet surface area on that day. Factor 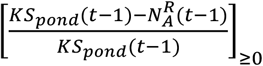 is non-negative.

Juveniles become adults at age *α*. Adults only die because of density dependence, or when the pond radius has become zero.

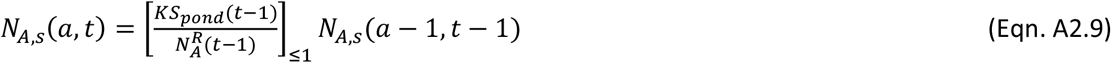

The density dependent factor 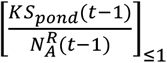 cannot be larger than one. Adult females which will survive until the next day *t* produce new eggs every day throughout their lives, which are fertilized by males, which will also survive until the next day. Reproduction requires that resident individuals of both sexes are present and it is also set at zero for very biased resident sex ratios (ratio resident females to males outside of the interval [*S*^-1^, *S*]). If all conditions are fulfilled, the number of new eggs with a mutant genotype appearing on day *t* equals

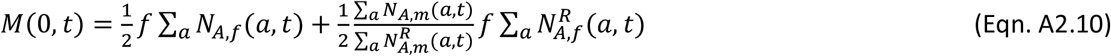

where the factors 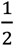 account for the fact that mutant individuals are heterozygotes and only transfer the mutant allele to half of their offspring. 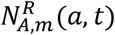 is the resident number of adult males and 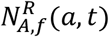 the number of resident adult females of a specific age present at time *t*. New eggs and their embryos have experienced zero dry days, they are put in wet conditions in the soil. These eggs are distributed at random over the pond bottom which adults can reach. This implies that location category *j* receives eggs proportional to its surface (which is *π*(*j* +)), giving the *M*_*j*_(0, *t*) per location category.

On the first day with dry conditions at location *j*, all new eggs deposited in that location in the preceding wet period *M*_*j*_(0, *t*) are added to the stage one embryos from older wet periods. The buffer with new eggs is emptied.

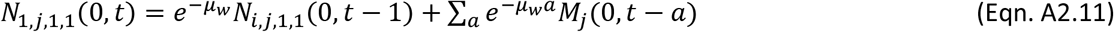

Resident individuals have the homozygous genotype and the phenotype of a homozygote. To determine their numbers we can use mostly the same equations as above where resident individuals are now counted. Only in the number of new eggs produced (Eqn. A2.10), a modification has to be made to make it represent the number of new eggs with resident genotype. Only the route via females needs to be accounted for and resident genotypes are transmitted with probability one, not one half as all males will contribute the resident allele when mutants are rare.

### Mutational effects on transition probabilities

Some more detail is required to understand what a mutation change in one of the rates will actually change to a transition matrix. If we make the elements of a minorized transition matrix ***M***_66_ from one day to the next explicit, we can write that matrix as consisting of transition probabilities *p*_*i*→*j*_ from state *i* to state *j*, where each state is a developmental stage (*i, j* in 1,…, 5).

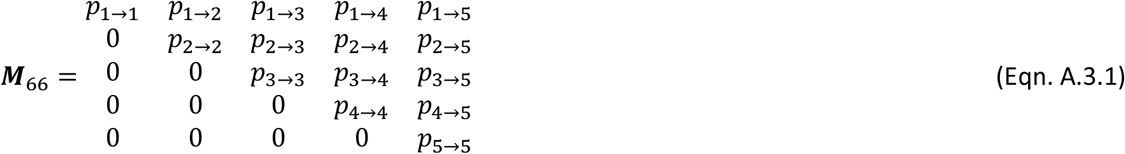

Transitions *p*_*i*→*dead*_ can be inferred from the other transitions as *p*_*i*→*dead*_ = − ∑_*j*≠*dead*_ *p*_*i*→*j*_. Diagonal elements in ***M***_66_ can be decomposed as *p*_*i*→*i*_ = − ∑_*j*≠*i*_ *p*_*i*→*j*_ − *p*_*i*→*dead*_. The probabilities of developmental transitions are not conditional on not dying. All transitions are treated on an equal footing. One needs to be careful when changing hazards to keep all elements within the bounds [0, 1] and to preserve the sum constraint on the probabilities out of a state. Therefore, when for a particular day, no events have been observed in the data (due to lack of events or of observations), the matrix for that day is not changed by trait changes. Also, when one of the probabilities on a row is one or the diagonal is zero, trait changes in the involved cumulative hazards are not affecting that row. Only diagonal elements *p*_*i*→*i*_ of ***M***_66_ (Eqn. A.3.1) with values within the interval ]0, 1[ are affected by trait changes to the cumulative hazards for death rates. It is assumed that trait changes in stage-specific rates of development, affect all cumulative hazards for changes out of that stage equally. For example, a multiplicative increase *c* in the rate of development out of stage one affects four cumulative hazards, changing all off-diagonal probabilities in ]0, 1[ in the first row of ***M***_66_ (Eqn. A.3.1) with a factor *c* and therefore reduces the diagonal with (*c* −) ∑_*j*=2,3,4,5_ *p*_1→*j*_. Trait changes studied were limited to the interval [0.9, 1.1] for which it was checked that all probabilities in transition matrices ***M***_66_ remain between zero and one.

### Invasion fitness

To determine invasion fitness and adaptedness, it is necessary to assume stationarity of the environment in the long run (Metz et al. 1992, Van Dooren 2022). If we want to assess whether organisms which are prevalent in a given environment are adapted to it, then we should determine in the first place that organisms with genotypes determining different but similar phenotypic strategies would not be able to increase in frequency when they would appear. This is the idea of adaptation bringing populations to a final stop of phenotypic and genetic evolution (Hammerstein 1996), which became formalized in adaptive dynamics approximations based on invasion fitness (Metz et al. 1992, Metz et al. 1996, Geritz et al. 1998). At such endpoints of evolution, invasion fitnesses of rare (mutant) phenotypic strategies slightly differing from the prevailing or so-called resident strategy should not indicate that these can invade. At adapted population states, the invasion fitness as a function of mutational changes is at an optimum and the first condition for that is that the linear approximation of such a function equals zero for small mutational changes, i.e., all partial derivatives (slopes) of invasion fitness with respect to a mutational change in a single trait are zero (Metz et al. 1996). When invasion fitness is estimated for mutants appearing in a resident population with stationary but non-constant numbers or densities, it is best to calculate initial and final numbers of mutants over a period which is a multiple of the periodicity in the resident population dynamics. Respecting this periodicity minimizes errors in estimation (Van Dooren & Metz 1998). All pond regimes investigated have a duration after which the same pattern of wet and dry periods repeats itself, and the resident attractors follow this periodicity. Invasion fitnesses were calculated from numbers at the start and end of a multiple of the total duration of the resident pattern.

As further supplementary material, two figures are provided.

**Supplementary figure one.**
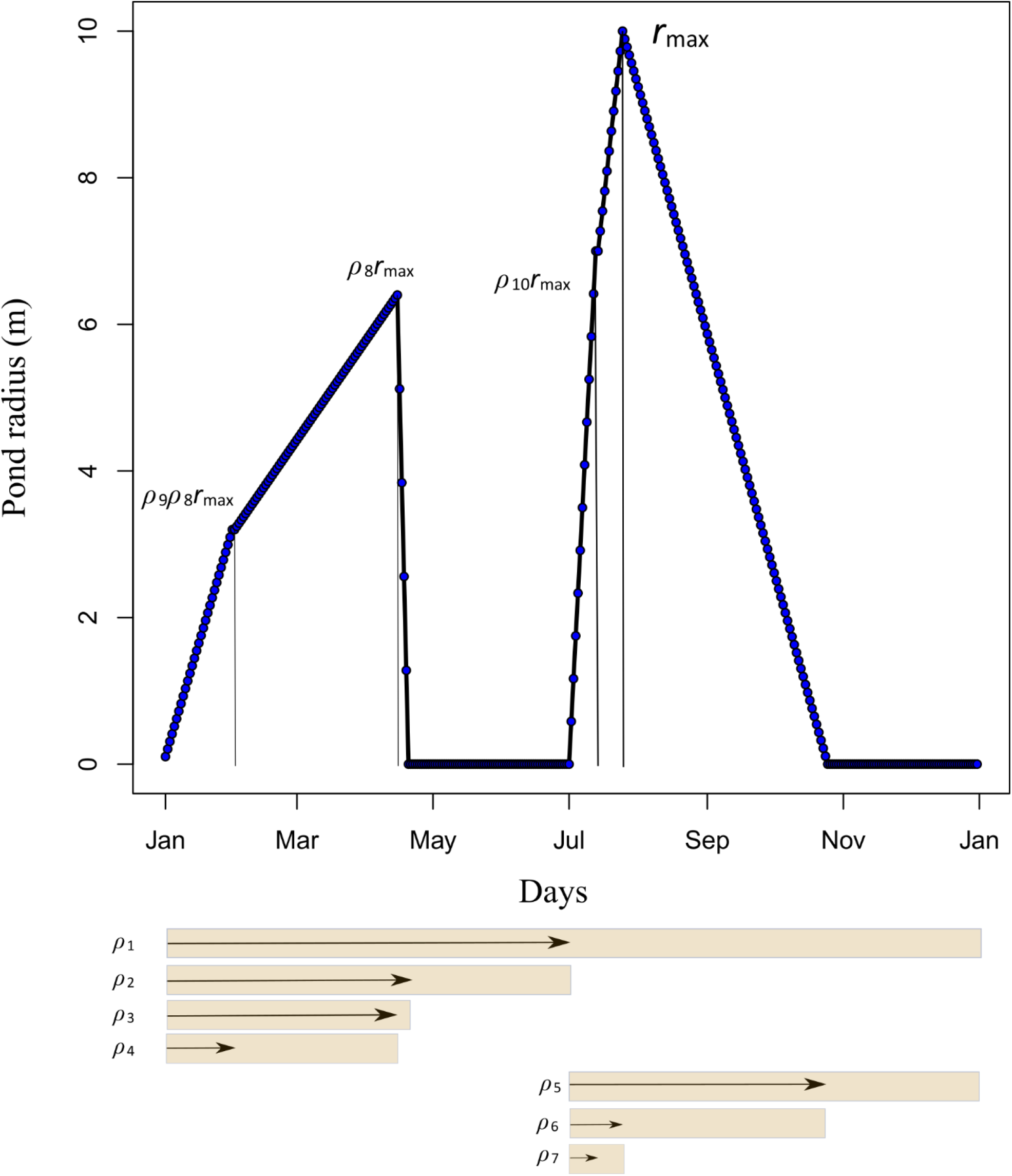
Parameterization of pond filling time profiles. Radiuses *r*_pond_(t) are piecewise linear continuous functions (when time is given as a real number). Differences between a large number of piecewise linear pond filling profiles can be specified by a change in one or several of ten parameters. These parameters, all constrained between zero and one, are involved in the specification of each pond filling regime, for a fixed parameter *r*_max_, the maximum radius the pond can have within a year. As they are all proportions, they are indicated in the figure by bars, which show the intervals on which each parameter works, and by arrows, showing the actual value they took to produce the pond profile shown. Parameter *ρ*_1_ is the proportion of the entire year taken up by a first pond filling and drying period. Parameter *ρ*_2_ is the proportion of that period which is wet. Parameter *ρ*_3_ if the proportion of that wet period at which maximum radius is reached. Parameter *ρ*_4_ is the proportion of the first wet period where the radius increases, at which the speed of filling changes. Parameters *ρ*_5_ to *ρ*_7_ are the corresponding parameters for the second pond filling and drying period. Parameter *ρ*_8_ is the proportion of *r*_max_ which is the maximum reached within the first period. Parameter *ρ*_9_ determines the radius at which the speed of filling changes in the first wet period and parameter *ρ*_10_ determines this radius for the second wet period.

**Supplementary figure two.**
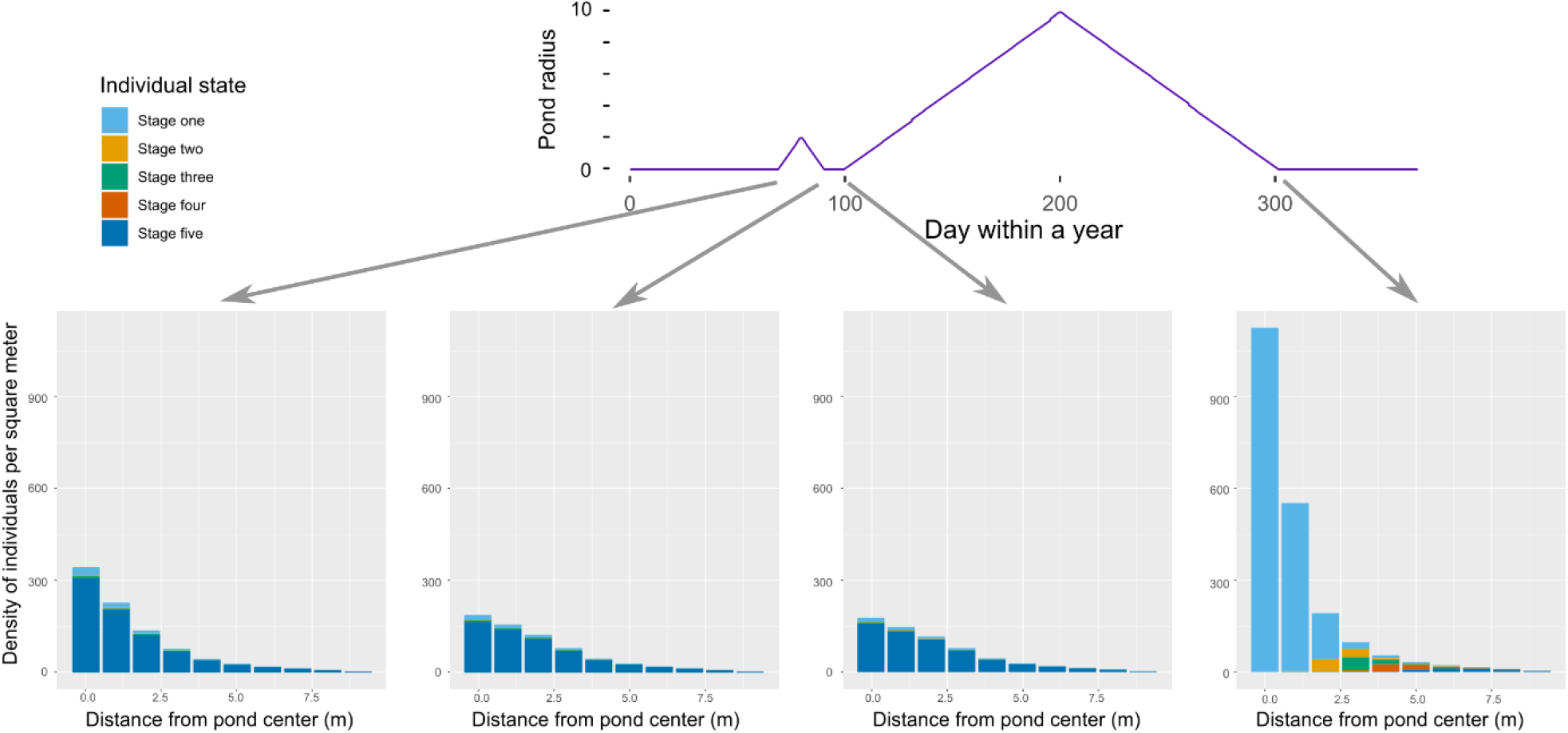
Distributions of embryos in different stages at different time points within a year. Distributions over stages at the end of each period in a pond filling regime drawn above the stacked barcharts are shown, as a function of distance to the pond center. The pond filling regime has a short period where reproductions cannot occur and a main longer one where new eggs are added to the egg bank. By the end of the wet period, on day 300, all eggs at nine to ten meter from the pond center have been dry since day 212.

## Notes

### Competing Interest Statement

The authors have declared no competing interest.

### Summary of Updates

The manuscript has been reviewed. Comments from three reviewers and an associate editor have been taken into account.

